# 5-Azacytidine: Effects on the Expression of alpha-Cardiac Actin in Pericytes from Human Adipose Tissue

**DOI:** 10.1101/2021.02.08.430216

**Authors:** Valéria Ferreira-Silva, Munira M. A. Baqui, Greice A. Molfetta, Aparecida M. Fontes, Dalila I. Zanette, Fernanda U. Ferreira, Maristela D. Orellana, Wilson A. Silva, Dimas T. Covas

**Affiliations:** Centre for Cell Therapy and Regional Blood Center, National Institute of Science and Technology in Stem Cell and Cell Therapy, Medical School, University of São Paulo, Ribeirão Preto, São Paulo, Brazil; Department of Cell and Molecular Biology and Pathogenic Bioagents, Medical School, University of São Paulo, Ribeirão Preto, São Paulo, Brazil; Department of Genetics, Medical School, University of São Paulo, Ribeirão Preto, São Paulo, Brazil; Department of Clinical Medicine, Medical School, University of São Paulo, Ribeirão Preto, São Paulo, Brazil

**Author notes:** Correspondence author (VF-S).

**Keywords:** 5-Azacytidine, DNA Methylation, Actin, Pericytes, Adipose-tissue derived stem cell, Cardiomyogenic differentiation, Epigenetic modification

## Abstract

DNA methylation patterns are closely related to the chromatin structure, and its remodeling is considered an important mechanism in the control of gene transcription during cell differentiation. In rodent, several studies have related the possibility that multipotent mesenchymal stromal cells (MSCs) undergo cardiomyogenesis. However, it has not been completely elucidated if human adult stem cell exhibits true differentiation potential for a cardiac lineage. In this study, the action of the DNA methylation inhibitor 5-azacytidine (5-aza) was examined in human adipose tissue pericytes (hATPCs: 3G5^+^) regarding their possible capacity to induce myocytes *in vitro*. Real-Time PCR revealed that cells treated with 5-aza presented time-dependent decrease in the mRNA expression of α-cardiac actin (α-CA). At 24 h, this diminution was statistically significant; however, there was not a correlation with the highest level of DNA demethylation at the same period using Methylation-Sensitive High Resolution Melting-PCR (MS-HRM-PCR). An evident increase in the α-CA protein expression was observed by Western blotting in hATPCs treated with 5-aza at 24 h. The mRNA expression of α-SMA (α-smooth actin) also showed a time-dependent decrease after the treatment, however, it was not significant. The ultrastructural analysis showed similar structures such as like-cell junctions, caveolae, and actin myofilaments, which aligned in parallel. These phenotypic alterations were found only after the treatment; however, the hTAPCs after 5-aza treatment were not able to form thick myofilaments and consequently sarcomeres. These results indicated that a terminal cardiac differentiation of hTAPCs was not achieved and that the cardiomyogenesis failure could be related to the non-muscle origin of the adipose tissue.

## Introduction

Multipotent mesenchymal stromal cells (MSCs), derived from human bone marrow, retain the ability to differentiate in some cell types *in vitro*, besides contributing to the regeneration of mesenchymal tissues such as bone, cartilage and adipose among others [24]. Adipose tissue, as well as bone marrow, are derived from the embryonic mesoderm and contains MSCs [32, 25, 16]. The procurement of this tissue in large quantities, using a minimally invasive method by liposuction, has conducted the researchers to consider it a good candidate for harvesting large numbers of MSCs for the studies on basic science and cell therapy [32].

Independent studies indicated that MSCs are located in a perivascular niche of different tissues [29, 9, 5, 32], and can be considered as reservoirs of multipotent progenitor cells. In addition, perivascular cells, including pericytes located in the microvessels and adventitial cells around large vessels [14] expressed MSCs markers showing similarities between both cell types [9, 5, 32]. Pericytes (also described as Rouget cells or mural cells and undifferentiated cells) are morphologically defined as cells situated around gap junctions, adjacent with endothelial cells in the microvasculature, and are continuous with the vascular basement lamina [14, 13]. Pericytes purified by fluorescence activated cell sorting showed to be myogenic in cell culture and *in vivo* after the isolation from skeletal muscle [11] and unexpectedly after the isolation of non-muscle tissues, such as human adipose tissue and pancreas [8].

To elucidate the mechanism involved in myogenic differentiation, there are several reports which describe *in vitro* assays to examine which cell population exhibits the highest potential to differentiate in myocytes [4, 11, 8, 22] and cardiomyocytes [19, 31, 7] or even which factors or pharmacological agents have influenced this differentiation. The DNA methylation inhibitor 5-aza was the first compound used to induce the formation of chrondrocytes, adipocytes and terminally differentiated striated muscle cells from fibroblasts in cultures originated from mouse embryo [4, 30]. Later, the mechanism of action of 5-aza was studied on stromal cells [19] or MSCs from bone marrow [34, 31], adipose tissue [7] and even on embryonic stem cells (P19) [6].

On the other hand, it is known that DNA demethylation induced by 5-aza alters chromatin structure, and this alteration is considered a determinant factor for epigenetic regulation of gene expression [20]. DNA methylation is a covalent modification in which the 5◻ position of cytosine is methylated due to a reaction catalyzed by DNA methyltransferases (DNMT1, DNMT3a, and DNMT3b) with the *S*-adenosyl-methionine (SAM) as the methyl group donor [21, 23]. In mammalian cells, this modification happens predominantly in CpG (cytosine-phosphate-guanine) dinucleotides, which are asymmetrically distributed in the genome, with CpG-poor regions and CpG-rich regions, also named CpG islands [3].

5-aza inhibits the DNMTs and, as a result, selectively activates genes silenced by methylation [20, 15]. Thus, it is believed that drugs which alter the DNA methylation pattern may induce cellular differentiation [4, 6, 3 15]. However, as far as we know studies evaluating the efficacy of 5-aza treatment on cardiomyogenesis of human pericytes have never been reported. Here, we evaluate *in vitro* the potential of hATPCs exposed to the 5-aza to induce cardiomyogenesis.

## Materials and Methods

### Ethics Statement

This study was approved by the Human Research Ethics Committee of the Clinical Hospital, Medical School of Ribeirão Preto, University of São Paulo. Written informed consent was obtained from each donor, in accordance with the Institutional Ethics Committee and with the 1964 Helsinki declaration and its later amendments or comparable ethical standards. A small fragment from right atrial appendage was obtained by coronary revascularization surgery (permit number, 12891/2010). Human fragments from uterus and stomach were donated from hysterectomy and gastric surgery samples, respectively (permit number, 12891/2010).

### Pericytes

Adipose tissue was acquired from elective liposuction procedures and cultured pericytes were provided by Dr. Lindolfo da Silva Meirelles, who developed a methodology for their isolation [10].

### Cell culture

hATPCs were cultured on poly-L Lysine (Sigma, St, Louis, USA) coated flasks (75 cm^2^, Greiner Bio-One, Frickenhausen, Germany) in specific medium for this cell type (ScienCell, Carlsbad, CA, EUA). After the confluence of approximately 70-85%, the cells were detached by treatment with Trypsin solution (0.025%) and EDTA (0.5 mM) (ScienCell, Carlsbad, CA, EUA) and aliquots of 1×10^5^ cells/well (6-well plates) were maintained in specific medium at 37°C and 5% of CO_2_. We performed cell expansion up to the fifth passage to obtain the sufficient number of cells for the cell differentiation experiments.

### 5-aza treatment

The cells were seeded in 6-well plates at a density 1×10^5^ cells/well. Twelve hours after seeding, the cells were washed with PBS twice and treated with 10 μM of 5-aza (Sigma, St. Louis, USA) in DMEM-F2 medium (DMEM-F2, Dulbecco’s modified Eagle medium: nutrient mixture F-12 (Gibco, Grand Island, EUA) containing 10% of characterized fetal bovine serum (FBS) (HyClone, Logan, UT, EUA), 100 U/mL of penicillin/streptomycin. The cell cultures were incubated at 37°C, 5% of CO_2_ and harvested after 2, 4, 24, 48, 72 h. Treatment extended up to one month.

### Morphological analysis

#### Phase-contrast microscopy

The cells were daily observed using inverted microscope (Olympus 1X71, Tokyo, Japan) by phase-contrast microscopy for morphological analysis. Photographic documentation was performed using a digital camera (Olympus UTVO 5XC-3, Tokyo, Japan).

### Transmission electron microscopy

1×10^5^ cells/well in 6-well culture plates were rinsed twice with PBS, fixed in 2% glutaraldehyde (Ladd Research Industries, Burlington, VT, USA) and 2% paraformaldehyde (EM Sciences, Fort Washington, PA, USA) in 0.1 M cacodylate buffer, pH 7.4, containing 0.05% CaCl_2_ for 1 h at room temperature. Cells were post-fixed in 1% OsO4 (EM Sciences, Fort Washington, PA, USA) in 0.1 M cacodylate buffer, pH 7.4, for 2 h, rinsed in ultrapure water, dehydrated in a graded series of ethanol. Cells were removed from the cell culture plates with propylene oxide and embedded in EMBED 812 (EM Sciences, Fort Washington, PA, USA). Thin sections were cut with a diamond knife, mounted on copper grids, and stained for 10 min each with Reynolds lead citrate and 0.5% aqueous uranyl acetate and examined in a Philips EM 208 transmission electron microscope.

### Real-time reverse transcriptase polymerase chain reaction

The isolation of RNA was performed with *RNeasy* micro kit (Qiagen, GmbH, Hilden, Germany) and the reaction of RNA (1 μg) reverse transcription to c-DNA using High-Capacity kit (Applied Biosystems, Foster City, CA, USA Boston, USA), following the protocol described by the manufacturer. Gene expression was assessed using TaqMan system (Applied Biosystems, Foster City, CA, USA) for a final volume of 15 μL in duplicates. The inventoried assay probes were α-AC (ACTC1, primer/probe Hs01109515_m1) and α-SMA (ACTA2, primer/probe Hs 00909449_m1). The constitutive gene glyceraldehyde-3-phosphate dehydrogenase (GAPDH_PN 4326317E) was used as an endogenous control. The reactions were processed in the equipment 7500 Real-Time ABI Prism Sequence Detection System (Applied Biosystems, Boston, USA) following the instrument’s default conditions. The results are presented as relative fold change in the gene expression, which was normalized to GAPDH gene. The baselines obtained from the control samples were used as a calibrator (non-treated hATPCs). The normalization and relative quantification of the gene expression were performed by the 2^-ΔΔ^C_T_ method, according to [17]. For the calibrator, ΔΔC_T_ was considered as zero and 2^0^ as one. For the samples treated with 5-aza, the value of 2^-ΔΔ^C_T_ shows the difference in fold in the gene expression related to the control.

### Identification of CpG islands

To identify possible CpG islands in *ACTC1* and *ACTA2* genes, DNA sequences were submitted to the online program EMBOSS http://www.ebi.ac.uk/tools/emboss/cpgplot/index.htlm

### MS-HRM-PCR

Genomic DNA was extracted from cells using the purification kit (Wizard, Promega, Madison, USA). Two μg of DNA was sodium bisulphite modified using the Eptech Kit (Qiagen GmbH, Hilden, Germany) in accordance with the manufacturer’s instructions. The reactions were conducted using the following reagents: 85 μL of sodium bisulphite, 2 μg of modified DNA, 35 μL of DNA protection buffer and nuclease-free water for a final volume of 140 μL. The reaction was gently mixed and transferred to Verity thermocycler (Applied Biosystems, Boston, USA), which was programmed to three denaturation cycles at 95°C for 5 min, followed by three incubation cycles at 60°C (25 min, 1 h and 25 min, and 2 h and 55 min) and a final cycle at 20°C. Afterwards, DNA was submitted to purification. The DNA amplification by MS-HRM-PCR was performed using: 10.0 μL of MeltDoctor MS-HRM Master Mix, 1.2 μL of each primer (5.0 μM), 2.5 μL of DNA modified, and nuclease-free water for a final volume of 20 μL. DNA varying between 0 to 100% of methylation were used as reference to build the standard curve. The reactions were kindly mixed and transferred to Veriti thermocycler (Applied Biosystems, Boston, USA). The amplification consisted of: 10 min at 95°C, followed by 40 cycles of denaturation 15 sec at 95°C and 1 min at the annealing. The melting curve was estimated in the following steps: one denaturation cycle of 10 s at 95°C, one annealing cycle of 1 min at 60°C, followed by one high resolution melting cycle (HRM) of 15 s at 95°C, and a last annealing step of 15 s at 60°C. MS-HRM-PCR analysis was done in *7500 FAST Real Time ABI Prism Sequence Detection System (*Applied Biosystems, Boston, EUA) using temperature ramp and acquisition of fluorescence in accordance with the manufacturers’ recommendation. Primers sequences used for bisulphite analyses were designed using Methprimer online program (http://www.urogene.org/methprimer/index1.htlm) and are set out in Table 1.

**Table 1.**
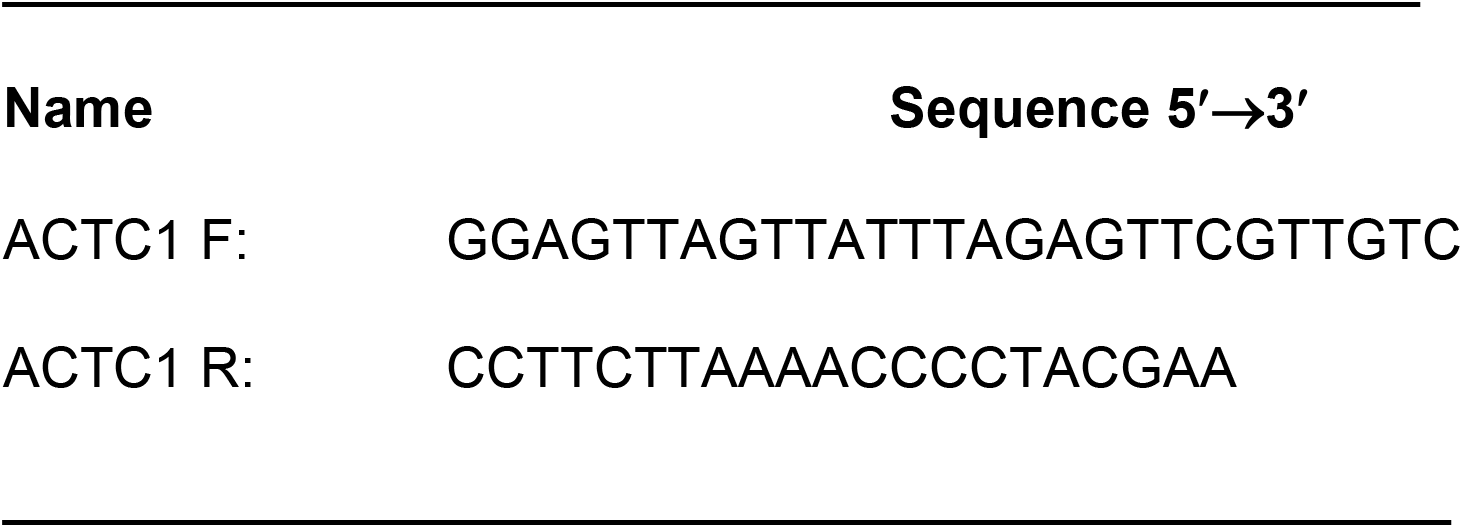
Primer sets used in the MS-HRM-PCR analysis.

### Western blotting

After harvesting, the muscle pieces were washed with DMEM (without FBS) and the tissues were kept on ice. Afterwards, the muscle were weighed and homogenized in Omni Tissue Homogenizer for 20-30 seconds using RIPA buffer containing protease and phosphatase inhibitor cocktail (Sigma, St. Louis, USA), centrifuged at 20.000 rpm for 20 min, at 4°C. Sample proteins were extracted using RIPA buffer and the concentration protein in the supernatant was determined by the Bradford method (Bio-Rad Lab, California, USA). Similar amount of total cell lysate were fractionated through a 10% SDS-PAGE and transferred to nitrocellulose membranes (Bio-Rad Lab, California, USA). The proteins transferred to the membrane were visualized by *Ponceau* solution stain 0.3% for 5 min. Later, the blots were blocked in TBS-T (50 mM Tris, pH 8.0, 150 mM NaCl, containing 0.1% Tween 20), containing 5% of nonfat dry milk. Expression of α-CA was detected using monoclonal antibody that recognizes the N-terminal region of the protein (clone AC1-20.4.2; Acris, Herford, Germany) and incubated with an anti-IgG mouse secondary antibody conjugated to peroxidase (Jackson ImunoResearch Laboratories, Inc, USA). The bands of the immunoreactive polypeptides were detected by chemiluminescence using ECL *plus* kit (Amersham Biosciences Co., Piscataway, NJ) according to the manufacturer’s instructions. Myofibril obtained from the skeletal muscles of rabbit was used as molecular mass standard [2].

Human smooth muscle from uterus and stomach were used as a control to confirm that α-CA is not present in these tissues.

### Statistical analysis

All values are expressed as the mean ± S.E.M. (n=3). Statistical analysis was carried with 2-way ANOVA followed by Bonferroni's post-test on GraphPad 5.0 Prism software (La Jolla, CA, USA). Values of *P* < 0.05 were considered significant.

## Results

### Phenotypic changes in hATPCs treated with 5-aza

The morphological analyses revealed that control hTAPCs, at all incubation times, kept fibroblast-shaped morphology (Fig 1a). After 5-aza treatment for 4 h, we observed a cell with an elongated tubular aspect, which seemed to have undergone successive nuclear divisions, showing various nuclei (apparently 4 or 5) within a common cytoplasm (Fig 1b).

**Fig 1.**
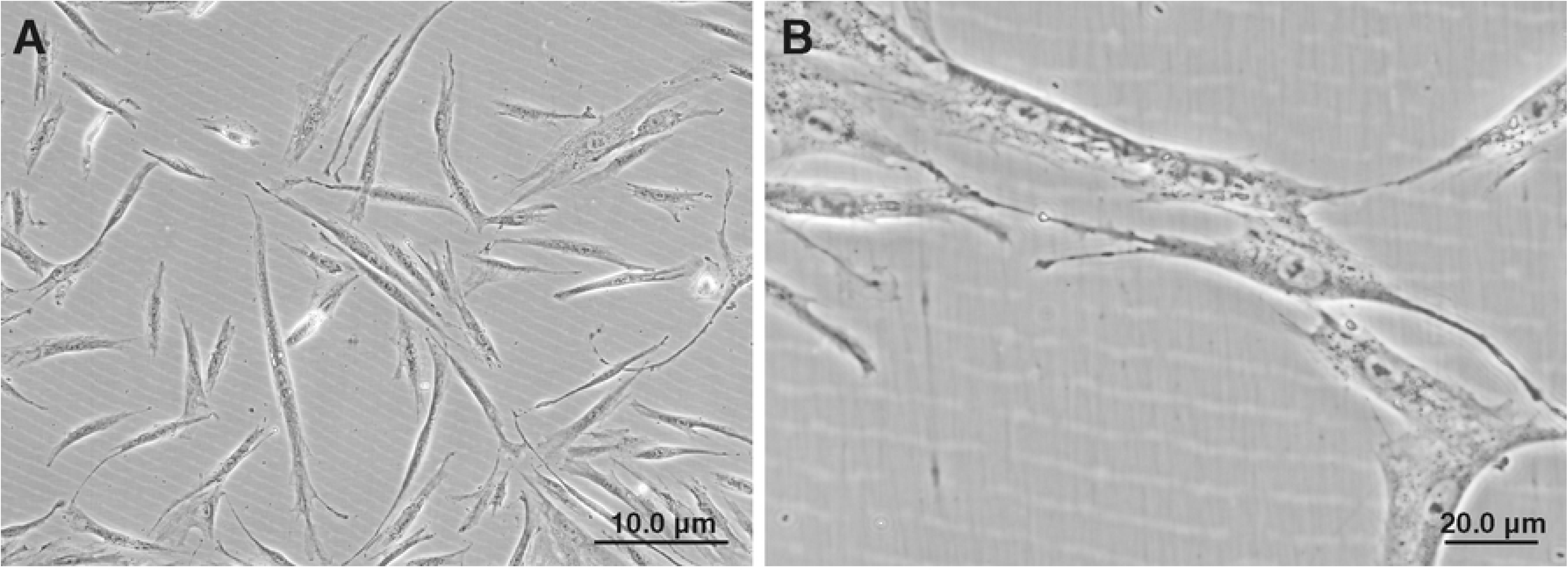
Phase-contrast morphology. Control hATPCs after 4 h of culture, showing fibroblast-like morphology (A). In 5-aza treated hATPCs for 4 h, we noticed a cell with an elongated tubular aspect, which seems to have undergone successive nuclear divisions, showing various nuclei (apparently 4 or 5) within a common cytoplasm (B).

The analyses by transmission electron microscopy revealed that 5-aza-treated for 4 h cells, but not the control cells (Fig 2a), exhibited similar structures to those found in muscles: like-cell junctions, caveolae, and the actin myofilaments in the cytoplasm aligned in parallel (Fig 2b). Actin myofilaments showed approximately a diameter of 5-9 nm, the standard size for this type of microfilament [1]. They were most highly concentrated just near the plasma membrane. On the contrary, control group showed the presence of microfilaments, but they were scarce and their alignment was less ordered (Fig 2a). However, the hTAPCs after 5-aza treatment were not able to form thick myofilaments and consequently sarcomeres. These results indicated that a terminal cardiac differentiation of hTAPCs was not achieved.

**Fig 2.**
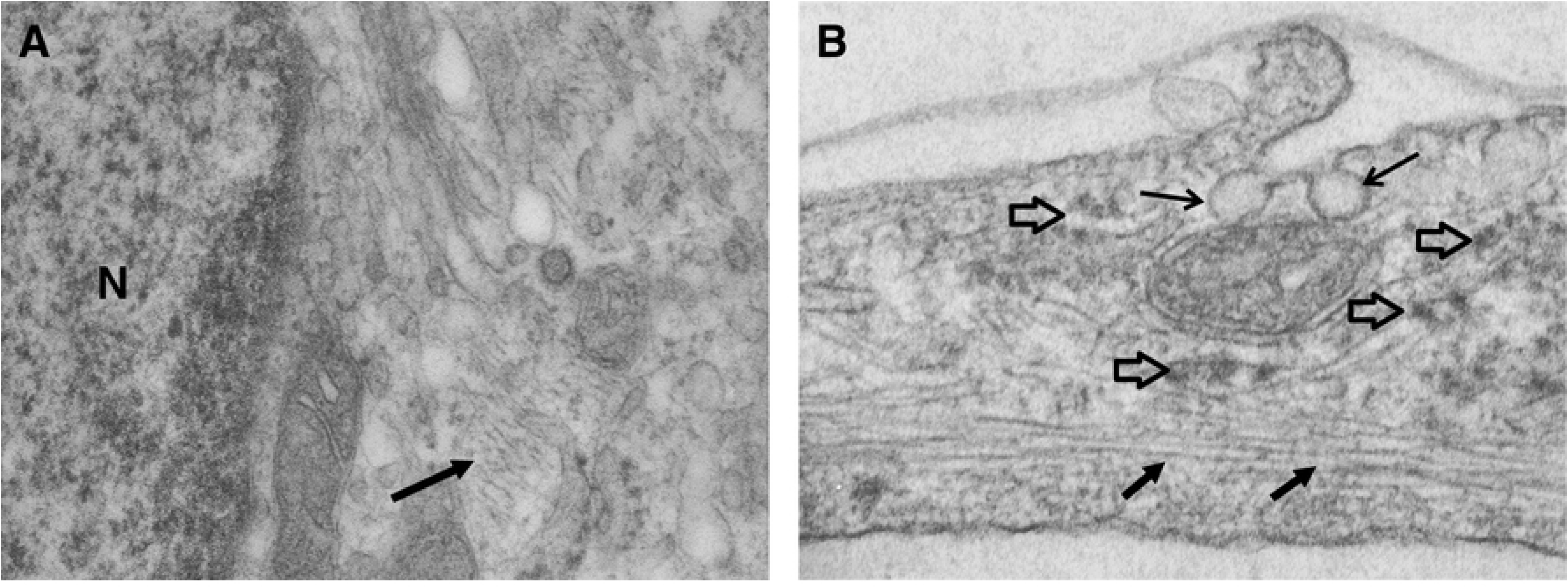
5-aza induced morphological alterations in hATPCs. Transmission electron micrographs from control hATPCs for 4 h in culture showed scarce microfilaments (broad black arrow) near the nucleus (N) (A). After 5-aza exposure for 4 h similar structures to those found in muscles: **like-** cell junctions (open arrows), caveolae on the cell surface (fine arrows), and actin myofilaments (broad bla**ck arrows) aligned in parallel, but** were not able to form thick myofilaments and consequently sarcomeres. **A**ctin myofilaments presented approximately a diameter of 5-9 nm, the standard size for this type of microfilament (B). The measurement was estimated using Image J software developed by National Institute of Health (NIH). (A, B: 50.000 x).

### Effects of the 5-aza on the α-cardiac actin mRNA expression

In the control hTAPCs, we observed α-CA mRNA basal expression at all-time points. 5-aza treatment induced a slight increase in α-CA mRNA expression in 1.27-fold in comparison with control group after 2 h (Fig 3). However, this increase has not showed any statistically significant difference. During the treatment, there was a time-dependent decrease in the expression levels from 0.85 to 0.31 between 4 and 24 h, respectively (24 h, statistically significant, *P* < 0.05) (Fig 3). Treatment after 24 h had no effect in α-CA mRNA expression.

**Fig 3.**
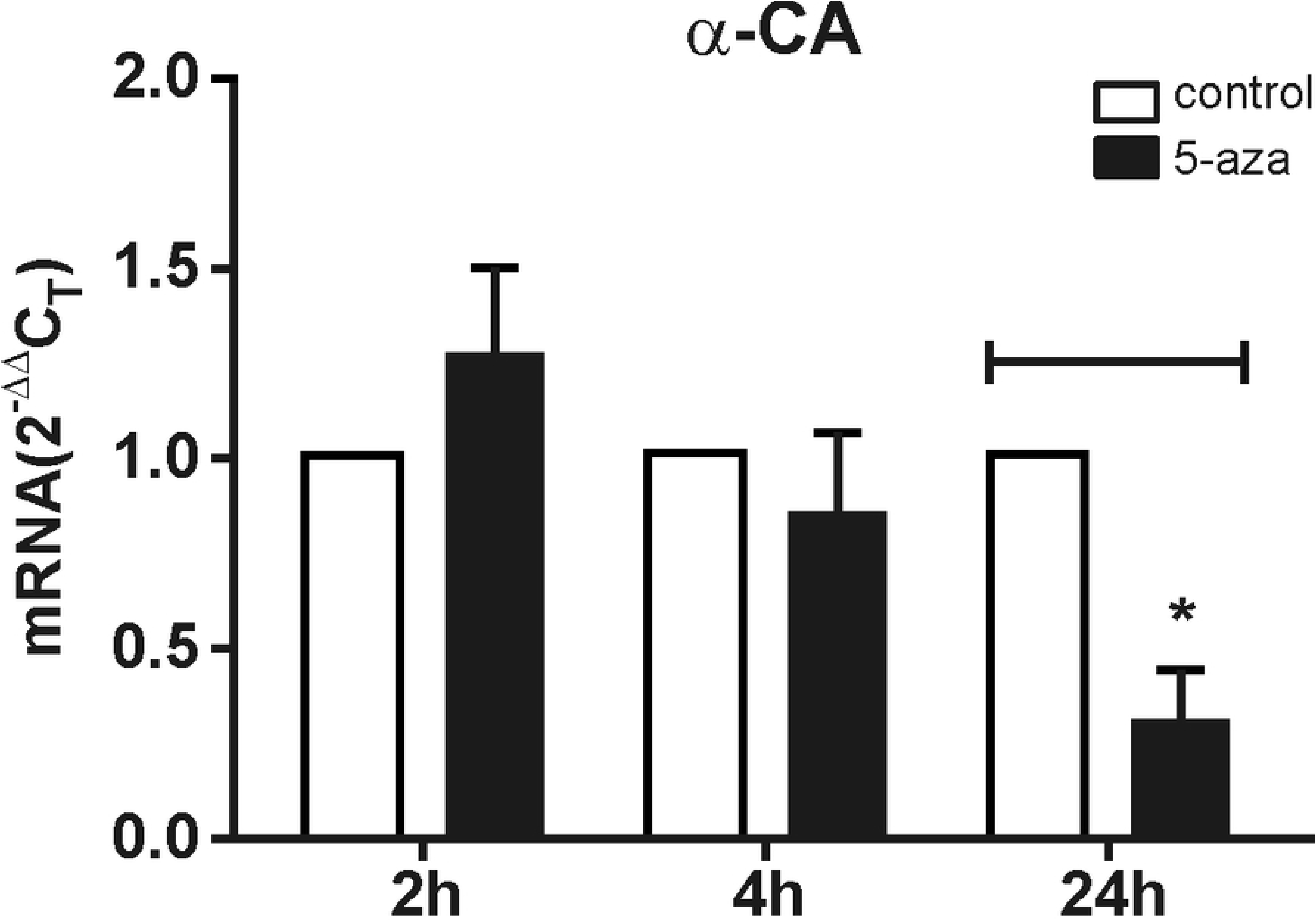
Effects of the 5-aza on the α-CA mRNA relative expression. hATPCs were cultured in the absence (control) or presence of 5-aza (10 μM) for different time periods. The results are expressed as 2^ΔΔ^C_T_, normalized to mRNA of GAPDH and calibrator (control hTAPCs). The bars show the mean ± S.E.M. of three independent experiments. (*) *P* < 0.05 in comparison with 5-aza treated cells, as evaluated by 2-way ANOVA, followed by Bonferroni’s post-test.

Levels of α-SMA transcript were detected in the control group at all-time points (Fig 4). 5-aza also caused time-dependent decrease in the α-SMA expression levels from 1.14-fold to 0.63 and 0.51 at 2, 4 and 24 h, respectively. However, at 24 h, the decrease of α-SMA expression was not statistically significant (*P*< 0.05) (Fig 4).

**Fig 4.**
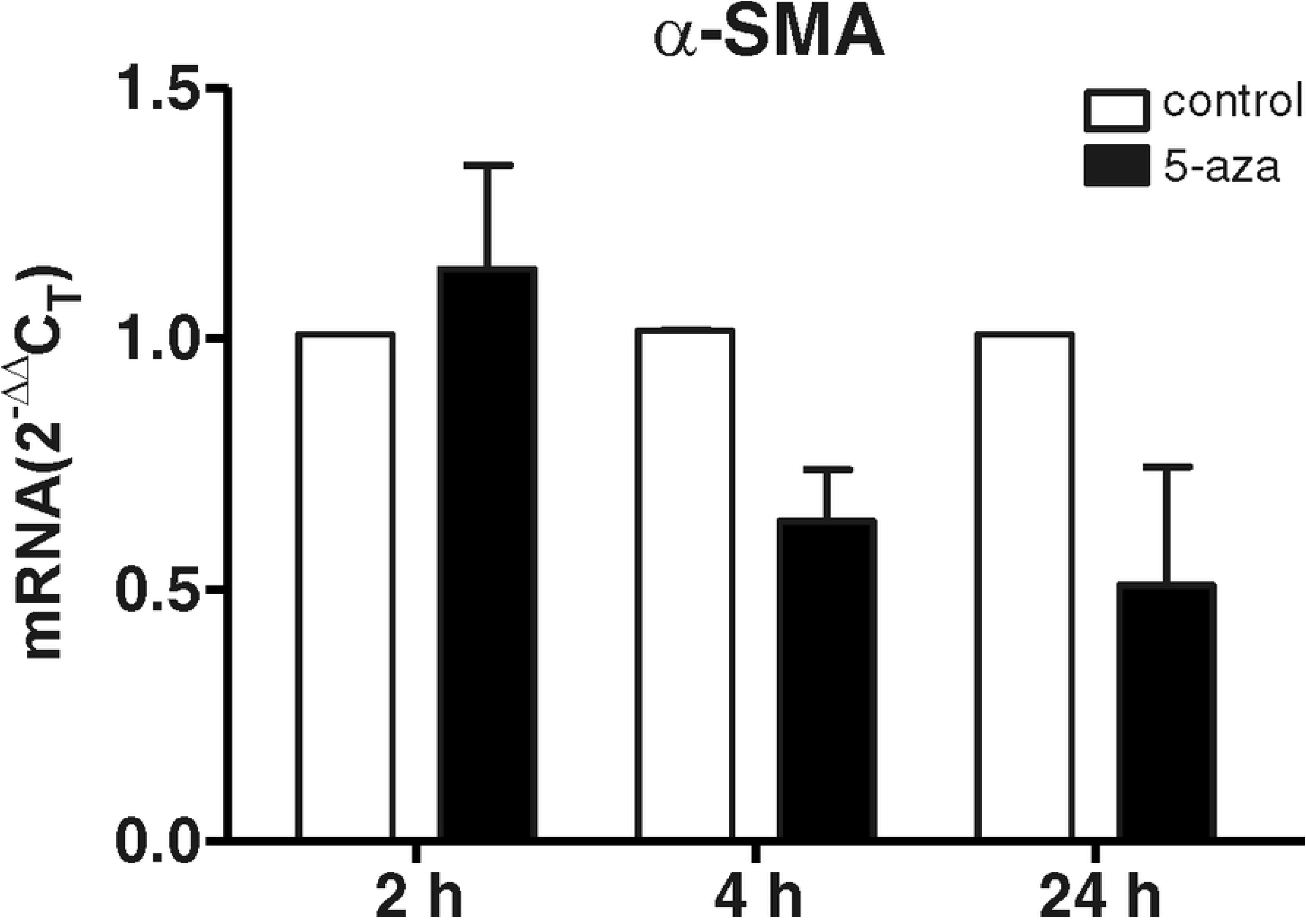
Effects of the 5-aza on the α-SMA mRNA relative expression. hATPCs were cultured in the absence (control) or presence of 5-aza (10 μM) for different time periods. The results are expressed as 2^ΔΔ^C_T_, normalized to mRNA of GAPDH and calibrator (control hATPCs). The bars show the mean ± S.E.M. of three independent experiments. Mean values of 5-aza treated cells were not significantly than that of control cells. *P* < 0.05, as evaluated by 2-*way* ANOVA, followed by Bonferroni’s post-test.

Untreated and 5-aza-treated hTAPCs have not expressed the markers of cardiac muscle: GATA-4 (GATA binding protein 4), NKx 2.5 (Homeobox protein NKx 2.5), β-MHC (β-Myosin Heavy Chain) and troponin T, besides of skeletal muscle markers Myf-5 (Myogenic regulatory factors-5) and Myo D (Myoblast Determination number 1) (used as differentiation controls) (data not shown).

### Effects of the 5-aza on the α-cardiac actin protein expression

α-Cardiac actin protein (43 kDa) was weakly detected by Western Blotting in control hTAPCs after 4 and 24 h (Fig 5a). Interestingly, in 5-aza-treated hTAPCs, we noticed an evident increase in α-CA expression after 24 h of treatment (Fig 5a).

**Fig 5.**
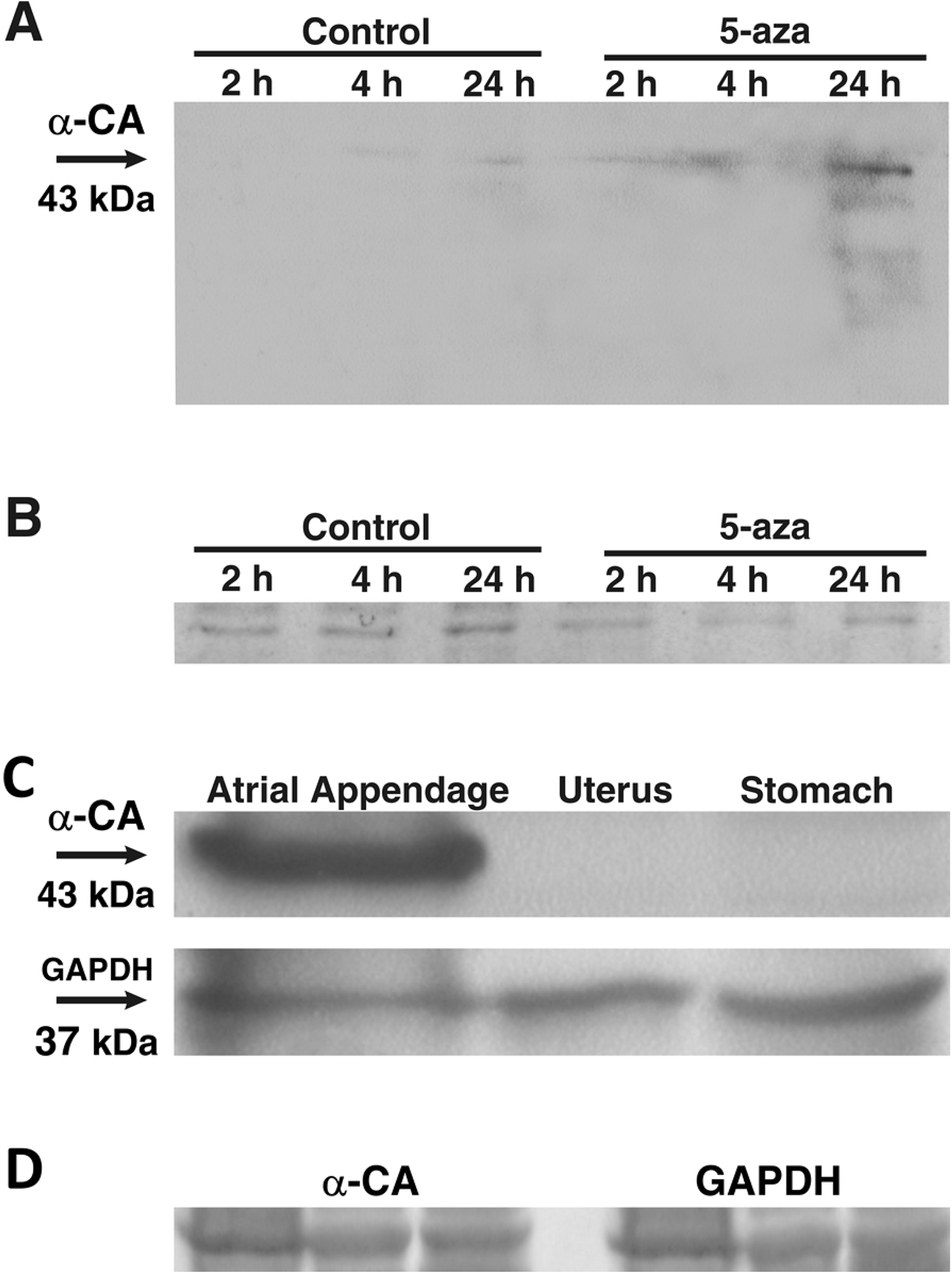
The alpha-CA protein expression of 43 kDa by Western blotting. After the treatment with 5-aza for 24 h, the 43 kDa band is clearly increased (A). Similar amounts of total proteins extracts were applied in each lane for separation, which can be visualized by staining of the membrane with *Ponceau* solution (B, D). The α-CA is strongly detectable in cardiac muscle (right atrial appendage), but is not detectable in smooth muscle (uterus and stomach) (C). Anti-GAPDH antibody was used as loading control (C).

To further confirm that the α-CA is not a constitutively expressed protein in other muscle tissues, but specifically expressed in cardiac muscle, we performed Western Blotting analyses using extracts of uterus, stomach (smooth muscle) and right atrial appendage (cardiac muscle). As expected, the results showed that α-CA is abundantly expressed only in heart (Fig 5c). Moreover, this experiment demonstrated the specificity of antibody that recognizes only the N-terminal region of the α-CA protein. Anti-GAPDH antibody was used as loading control (Fig 5c).

Similar amounts of total proteins extracts were applied in each lane for separation, which can be visualized by staining of the membrane with *Ponceau* solution (Fig 5b, c).

### Identification of CpG island in*ACTC1* gene (α-AC)

The analyses revealed the presence of one CpG island located in the promoter region of *ACTC1* gene (NM_005159.4) between 0.2 and 0.4 (x 10^4^) bp. This region corresponds to −562 and −770 bp, position given from transcriptional start site in the promoter region of *ACTC1* (Fig 6). On the contrary, no CpG island was found in the promoter region of *ACTA2* (NM_001141945.1) (data not shown).

**Fig 6.**
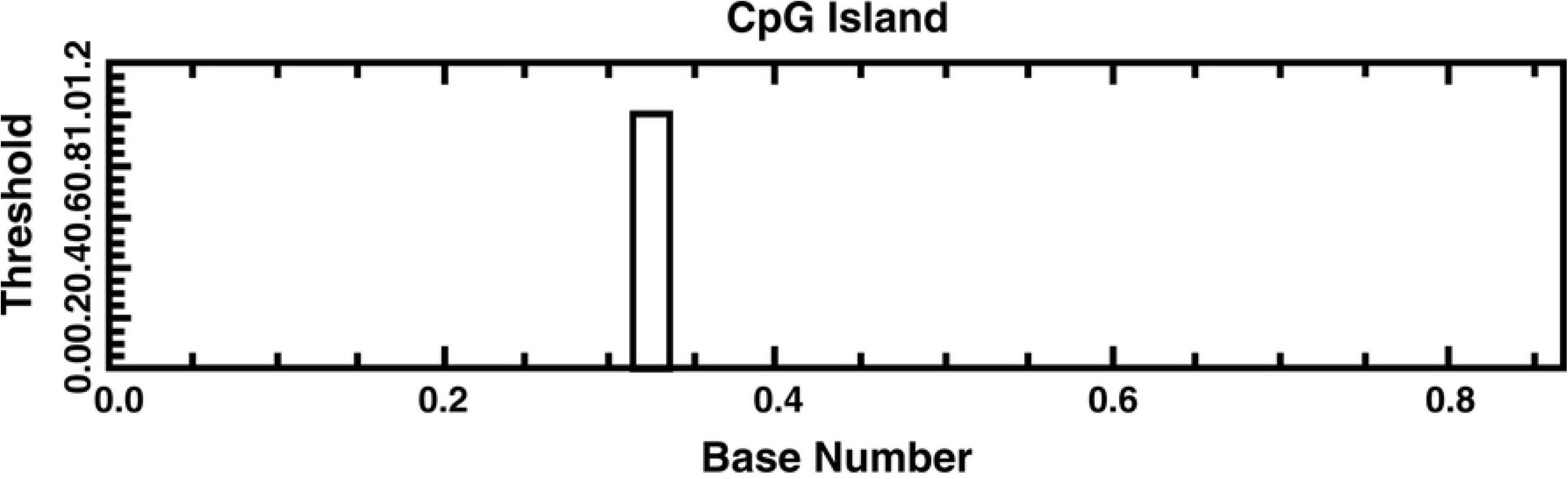
Location of one CpG island between 0.2 and 0.4 (x 10^4^) bp. This region corresponds to −562 and −770 bp, position given from transcriptional start site in the promoter region of *ACTC1* gene (α-CA). The sequence of *ACTC1* gene (NM_005159.4) was scanned by the online program: EMBOSS CpGPlot/CpGReport/Isochore.

### 5-aza-induced DNA methylation inhibition

Standard DNA varying between 0 to 100% of methylation (Fig 7a) was used to compare the DNA dissociation curves corresponding to CpG island of *ACTC1* gene promoter region. The results revealed that control hTAPCs presented around 90% of DNA methylation after 2, 4 and 24 h of cell culture (C2, C4 and C24, respectively) (Fig 7b). After 5-aza treatment, the level of DNA methylation was approximately: 80% at 2 h (A2), 40% at 4 h (A4) and 19% at 24 h (A24) *versus* 90% for control cells in all time (Fig 7b). These data showed the efficacy of the drug in the inhibition of methylation, and its time-dependent action mechanism in hTAPCs.

**Fig 7.**
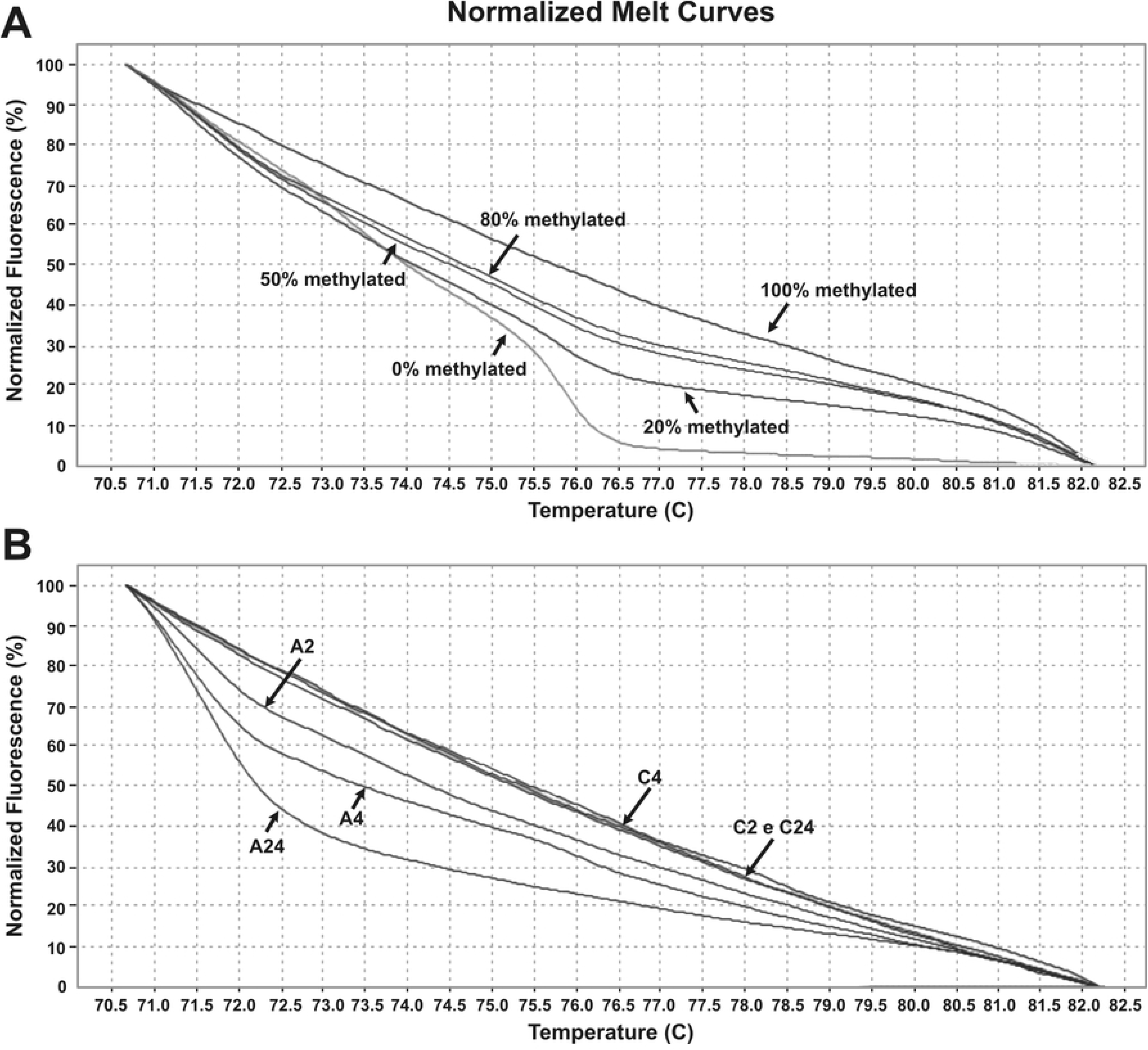
5-aza-induced DNA methylation inhibition in *ACTC1* gene by MS-HRM-PCR. Double-strand DNA labeled with fluorescent compound (*MeltDoctor*) was subjected to high temperatures (axis x), inducing DNA denaturation and, in consequence, the loss of fluorescence (axis y), generating different profiles of dissociation curves. Standard DNA ranging between 0 to 100% of methylation (A) was used to compare then with DNA dissociation curves corresponding to CpG island of *ACTC1* gene (α-CA) (B).

## Discussion

5-aza is routinely used as a chemotherapeutic agent against myeloid leukemia among other disorders [33] and has been extensively used for epigenetic research.

Generally, DNA demethylation induced by the drug would be responsible for the transcriptional activation of genes previously silenced which could be linked to cell differentiation [20, 15, 26]. Although cellular differentiation is usually unidirectional *in vivo*, this process can be reprogrammed and reverted *in vitro* [3].

The expression of α-SMA is commonly detected in pericytes from different tissues, including non-muscle tissues, and its regulation is extensively studied as a way of understanding the mechanism of cell differentiation [27, 28]. However, little is known about the regulatory events which govern α-cardiac actin (*ACTC1/*α-CA) expression after 5-aza treatment.

In the present study, bioinformatics data revealed the presence of one CpG island in the *ACTC1* promoter region. The analysis by HRM-PCR showed that after the treatment with 5-aza there was inhibition of time-dependent DNA methylation. At 24 h, 5-aza induced the highest level of inhibition (only 19% of DNA methylated relative to that in the control cells, 90%), showing an inverse correlation with the results obtained from the mRNA expression of α-CA. On the other hand, at the same time (24 h), there was an evident increase in the expression level of α-CA protein.

One possible interpretation of the results presented herein is that the basal levels of α-CA mRNA expression in control cells suggest that the gene is not fully repressed. Consequently, these basal levels could be limiting a relevant expression of the gene. Consistent with this Choi et al. (2010) also showed a time-dependent decrease in the α-CA mRNA expression during 5-aza treatment and inability to differentiate into cardiac lineage in MSCs from adipose tissue [7]. There is evidence that 5-aza could induce DNA global demethylation patterns, which could be influencing in the expression profile, but this mechanism is not yet completely elucidated [23].

Additionally, Lucarelli et al. (2001) related a strong correlation between the demethylation level of the sole CpG site and myogenin (myogenic differentiation factor) expression in mouse muscle cell line (C2C12) [18].

Our study showed that no CpG island was found in the α*-SMA* promoter region and there was no significant difference in the mRNA expression profile in different times of the treatment. As far as we known, there is no publication correlating the absence or presence of CpG island in the α*-SMA* promoter region with its transcriptional regulation in human cells, but only in rodent cells. It is believed that the extent and pattern of genomic DNA methylation is species and tissue-specific [20, 15]. In rats, it was observed that α*-SMA* gene has three CpG islands residing in the promoter and the first intron in fibroblasts with the capacity for myofibroblast differentiation [15]. In these cells, the findings revealed a distinct methylation pattern: more than 78% methylation in the promoter sequences, and less than 8% in the intronic region. This result was in contrast to lung alveolar epithelial cells that do not express α-SMA, which showed uniformly high methylation in comparable sequences of the α-SMA gene [15]. Thus, consistent with its role in gene silencing, highly methylated DNA sequences in the α-SMA gene correlated with suppression of its expression gene. Moreover, the effect of 5-aza on α-SMA expression was examined in fibroblast, showing a direct correlation between the increase of mRNA and protein expression. Therefore, inhibition of DNA methylation by the drug resulted in the induction of α-SMA expression during myofibroblast differentiation [15].

Although the primary amino acid sequence of actin has remained highly conserved during evolution, biochemical heterogeneity has been shown in the seventeen-amino terminal amino acids of actins from different tissues [27]. There are six actin isoforms produced by different genes in mammalian cells: two are located in virtually all cell types, and are named β-and α-actin; two are present practically in all smooth muscle cells (SMC), and are named α- and α-SM actin, and the other two are specifically expressed in striated muscle cells, and then referred to as α-skeletal and α-cardiac actins [27]. The analysis of cytoskeletal proteins has provided means to assess the differentiation state and the embryonic derivation of many cell types. The demonstration that pericytes from capillaries of rat pancreas expressed α-SMA was very important. The authors noticed that, besides pancreas, pericytes of the aorta and cardiac and skeletal muscles also expressed the marker [28]. Thus, the observation of muscle and non-muscle actins in pericytes from many organs supports the hypothesis that pericytes, at least in part, are multipotent [14, 12]. In accordance, da Silva Meirelles et al. [9] observed a similar pattern of expression for α-SMA among MSCs from different organs, indicating the existence of similarities between MSCs and pericytes.

More recently, Dellavalle et al. [11] described that pericytes from adult human skeletal muscle would be a second myogenic precursor (distinct from satellite cells), with myogenic potential to differentiation observed in culture. In this case, pericytes were also able to generate myofibres in dystrophic mice [11]. Despite this, Crisan et al. [8] reported that human perivascular cells, including pericytes, have innate myogenic potential independent on the origin tissue, such as the cells derived from the adipose tissue, pancreas, placenta, and skeletal muscle. The muscle regeneration index of fat-derived perivascular cells was even higher than of muscle perivascular cells [8].

In the present investigation, no evidence of the cardiac markers expression: NKx2.5, GATA-4, β-MHC and troponin-T, as well as of the early markers of skeletal muscle differentiation pathway (Myf-5 and MyoD) was observed in control and 5-aza-treated hTAPCs.

Taken together, the results indicated that the increase expression level of α-CA protein induced by 5-aza treatment is insufficient for the triggering of cardiomyogenesis *in vitro*, and that the failure could be related to the non-muscle origin of the adipose tissue.

## Acknowledgments

Special thanks to Dr. Lindolfo da Silva Meirelles by generously providing the pericytes. Dr. Solange Basseto, Dr. Mauricio M. Sabino de Freitas (*in memoriam*) and Dr. Alberto F. Gaspar (University of São Paulo) and their patients for tissues procurement (right atrial appendage, uterus and stomach respectively). Sandra N. Bresciani for artwork and Fernanda Udinal for language advice.

## Author Contributions

**Conceived and designed the experiments**: VFS MMAB GAM DLZ

**Performed the experiments:** VFS MMAB FUM GAM DLZ

**Analyzed the data**: VFS MMAB FUF DLZ GAM

**Contributed reagents/materials/analysis tools:** MDO AMF WAS jr. DTC

**Wrote the paper:** VFS

**Supervision:** DTC

